# SLIMP: Supervised learning of metabolite-protein interactions from co-fractionation mass spectrometry data

**DOI:** 10.1101/2021.06.16.448636

**Authors:** Boris M. Zühlke, Ewelina M. Sokolowska, Marcin Luzarowski, Dennis Schlossarek, Monika Chodasiewicz, Ewa Leniak, Aleksandra Skirycz, Zoran Nikoloski

## Abstract

Metabolite-protein interactions affect and shape diverse cellular processes. Yet, despite advances, approaches for identifying metabolite-protein interactions at a genome-wide scale are lacking. Here we present an approach termed SLIMP that predicts metabolite-protein interactions using supervised machine learning on features engineered from metabolic and proteomic profiles from a co-fractionation mass spectrometry-based technique. By applying SLIMP with gold standards, assembled from public databases, along with metabolic and proteomic data sets from multiple conditions and growth stages we predicted over 9,000 and 20,000 metabolite-protein interactions for *Saccharomyces cerevisiae* and *Arabidopsis thaliana*, respectively. Extensive comparative analyses corroborated the quality of the predictions from SLIMP with respect to widely-used performance measures (e.g. F1-score exceeding 0.8). SLIMP predicted novel targets of 2’, 3’ cyclic nucleotides and dipeptides, which we analysed comparatively between the two organisms. Finally, predicted interactions for the dipeptide Tyr-Asp in Arabidopsis and the dipeptide Ser-Leu in yeast were independently validated, opening the possibility for future applications of supervised machine learning approaches in this area of systems biology.

## Introduction

Small-molecules, whether a metabolite, short peptide, or drug, bind to a macromolecular partner, most commonly a protein, to exert their function (X. S. Li 2011). Binding can be both non-covalent or covalent and is associated with a conformational change of the protein target that results in a downstream effect. Small-molecule regulation of proteins, ranging from enzymes and transporters to transcription factors and structural proteins, constitutes an evolutionary conserved mechanism that enables organisms to respond to the developmental and environmental cues (M. S. Kosmacz 2020). While covalent modification often requires additional enzymes (e.g. (de)acetylase), non-covalent interactions are determined by the binding affinity, enabling fast and reversible regulation driven by the change in metabolite concentrations. The best-studied example of a metabolite-protein interaction (MPI) network is that of competitive or allosteric regulation of enzyme activities by small-molecules that share similarity with either the substrate or the product of the enzyme (Link 2013, Reznik 2017, Alam 2017, M. N. Diether 2019, Razaghi-Moghadam 2021). However, it has become increasingly clear that similarly to enzymes, many more proteins can bind and be regulated by a plethora of small-molecule ligands (Lempp 2019, Gallego 2010, X. G. Li 2010, Piazza 2018).

Despite their importance, cell-wide characterisation of the protein-metabolite complexes has lagged behind protein-protein interaction (PPI) studies, limited by the lack of suitable experimental approaches. Emergence of novel methods to capture MPIs and improvements in mass-spectrometry proteomics and metabolomics technologies now allow the use of the synthesized knowledge along with computational approaches to discover new MPIs (M. S. Luzarowski 2019, M. S. Diether 2017). We have recently adapted the co-fractionation mass spectrometry (CF-MS) method, historically used for protein-protein (McWhite 2020) and protein-RNA (Mallam 2019) interaction studies, for cell-wide characterization of MPIs (D. S. Veyel 2018, D. K. Veyel 2017), in an approach referred to as PROMIS. PROMIS relies on the size-based separation of molecular complexes present in the native lysate. While metabolites present in the protein complexes separate into high molecular weight fractions, free small-molecules separate in the slower eluting fractions of low molecular weight. Collected fractions are subjected to concurrent protein and metabolite extraction, followed by proteomic and metabolomic mass spectrometry-based analysis. Protein and metabolite elution profiles are in turn deconvoluted into single peaks, and co-elution is used to delineate putative protein-protein and metabolite-protein interactors. What distinguishes PROMIS from other methods used to study ligand-protein complexes is that instead of focusing on a single compound, it enables untargeted, metabolome-wide identification of metabolite-protein complexes. However, and as the separation is limited to 30-40 fractions, on average, every metabolite will co-elute with several hundred of proteins, of which only one may constitute a true interactor, while the remaining ones represent coincidental co-elution. To identify *bona fide* MPIs we complemented PROMIS with the orthogonal small molecule centred approaches such as affinity purification and thermal proteome profiling (M. V. Luzarowski 2021, D. S. Veyel 2018, M. L.-B. Kosmacz 2018). While the latter two approaches are limited to investigating MPIs that include known metabolites, available as purified compounds, and are highly laborious and time-consuming, the main advantage of PROMIS is its untargeted exploration of MPIs.

Here we explored an alternative approach to identify MPIs by using the established knowledge of interactions between metabolites and proteins present in our data sets, in a form of a gold standard, in combination with supervised machine learning methods. We hypothesize that, unlike coincidental, true MPIs will be predicted as present across different CF-MS data sets within and across species. Notably, combination of multiple CF-MS data sets has been successfully employed to identify functional protein-protein interactions (Wan 2013). To test our hypothesis, we combined multiple PROMIS experiments corresponding to different growth stages for two model organisms, the plant *Arabidopsis thaliana* and budding yeast *Saccharomyces cerevisiae.* We refer to the combination of CF-MS data sets with supervised machine learning method as SLIMP, for **S**upervised **L**earning of **I**nteractions between **M**etabolites and **P**roteins. As a result, we predict 9,434 MPIs between 54 metabolites and 1,556 proteins in *Saccharomyces cerevisiae* and 20,148 MPIs between 78 metabolites and 4,202 proteins in *Arabidopsis thaliana*. Extensive analyses corroborated the quality of the predictions from SLIMP with respect to widely-used performance measures (e.g. F1-score exceeding 0.8) and characteristics (e.g. learning and ROC curves). Functional characterisation of predictions from SLIMP verified known and provided a list of previously unknown MPIs. In summary, we demonstrate that the combination of multiple PROMIS experiments with SLIMP can be used to get insights in the complexity of metabolite-protein interactome across multiple species.

## Results and Discussion

### Data sets employed in prediction of metabolite-protein interactions

To predict metabolite-protein interactions we relied on proteomic and metabolomic profiles obtained from size exclusion chromatography (SEC) followed by liquid chromatography - mass spectrometry (LC-MS) applied on cell lysates from experiments with: (i) three different data sets from *Arabidopsis thaliana* as well as (ii) four different data sets from *Saccharomyces cerevisiae* (Supplemental Table S1). If a protein and a metabolite form a complex, these molecules are expected to co-elute and, hence, are found in the same fractions (D. S. Veyel 2018, M. V. Luzarowski 2021). Unbound metabolites are much smaller than proteins and elute last, making fractions separable into those that contain only metabolites and those that include both proteins and metabolites. For the performed experiments, 36-40 SEC fractions that include both proteins and metabolites were collected, covering sizes from 13 to 6037.7 kDa (see Methods). To increase the quality of the predictions, the data from experiments on each organism were jointly employed for feature engineering. As a result, we used 31% of all proteins and 24% of all metabolites, as they were detected over all experiments in *A. thaliana* and 36% and 17% of proteins and metabolites, respectively, in *S. cerevisiae* experiments (Supplemental Table S1). Further, the metabolomic and proteomic profiles from different data sets and replicates were combined (Methods). The pooling of elution profiles from different replicates of one experiment is justified if they are concordant. Indeed, we found that the metabolomic and proteomic profiles over the different fractions were highly concordant, as quantified by the R_V_-coefficient of value at least 0.7 (Figure 1A) (Abdi 2007).

**Figure 1:**
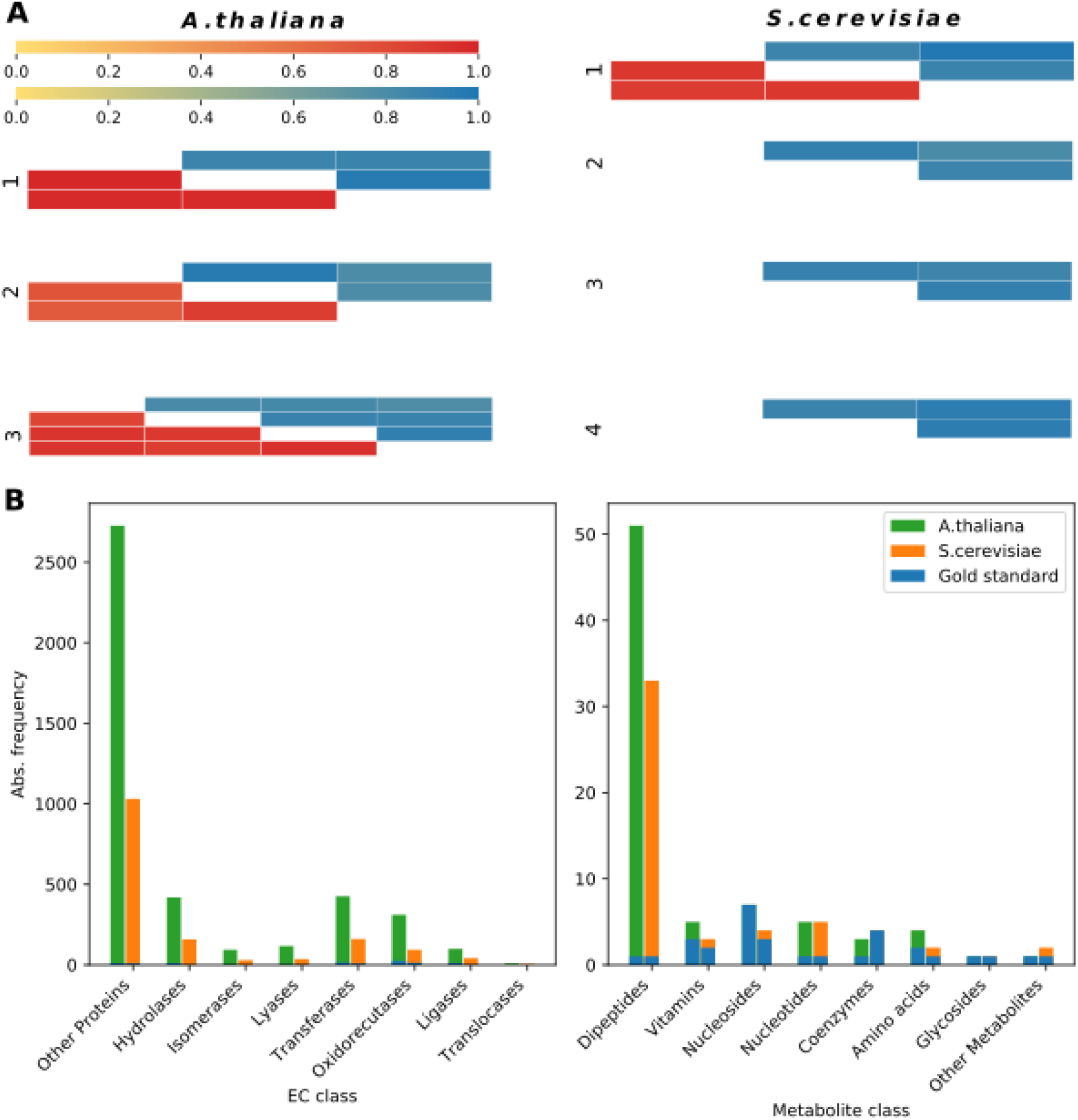
Descriptive analysis of the data sets employed in SLIMP. (**A**) Experimental reproducibility. Heatmaps describe the R_V_-coefficients of proteomic (red) and metabolomic (blue) profiles over the replicates of each experiment. (**B**) Protein and metabolite class distributions. Histograms present the number of proteins per assigned EC number (left) (Webb 1992) and metabolites per KEGG BRITE metabolite class (right) (Aoki-Kinoshita 2007) in the performed experiments on *A. thaliana* (green), *S. cerevisiae* (orange) and the corresponding training sets (blue).

We used BRITE classes and KEGG pathways (Aoki-Kinoshita 2007) to annotate the proteins and metabolites; moreover, proteins were also classified with enzyme commission numbers (Webb 1992) and GO terms (Gene Ontology Consortium 2008) (Supplemental Table S2). The used annotations showed that the data sets from *A. thaliana* and *S. cerevisiae* included proteins and metabolites from various processes, pathways and classes, indicating that they can be used to get a broad overview of MPIs (Figure 1B).

### Generation of a gold standard for supervised learning of metabolite-protein interactions

Building a supervised approach to learn metabolite-protein interactions requires access to a high-quality gold standard (Hastie 2009). We collected information about presence and absence of interactions from the STITCH (D. S. Szklarczyk 2016) and PubChem (Y. S. Wang 2013) databases to assemble positive and negative gold standards for *A. thaliana* and *S. cerevisiae*, respectively. The gold standards contain more proteins than metabolites, as is the case in the employed data sets (Supplemental Table S1). Moreover, we ensured that the gold standards are balanced with respect to the number of interacting and non-interacting metabolite-protein pairs, referred to as positive and negative instances (Supplemental Table S3). While the positive and negative gold standards are balanced, we noted that they cover 22 % and 26 % of metabolites, 2% of proteins and 0.04 % and 0.06 % of MPIs occurring in the intersection over the gathered experimental datasets for *A. thaliana* and *S. cerevisiae*, respectively. Nevertheless, the assembled gold standards provided a good coverage of metabolite and protein classes from the abovementioned annotations (Figure 1B).

### Feature engineering for predicting metabolite-protein interactions in SLIMP

The elution profiles were normalized to their profile’s maximum peak intensity and combined over experiment replicates using a selected pooling method, such that for every metabolite and protein a data profile per experiment was constructed (Methods). Next, we combined the resulting proteomic and metabolomic data profiles to arrive at a feature vector employed in machine learning. To this end, for every metabolite-protein pair we calculated the cross-correlation profile whose elements correspond to the sum of fraction-wise products of protein and metabolite elution profiles at a certain displacement (Figure 2, Methods). The cross-correlation profile of co-eluting molecules, that are assumed to interact, has a peak at zero displacement (Figure 2A). In contrast, proteins and metabolites that do not co-elute show no peak at zero displacement (Figure 2B). Cross-correlation peaks at medium displacement may indicate that the metabolite interacts with another protein eluting close to the investigated protein. We assumed that larger displacements do not hold information about the interaction between the investigated metabolite-protein pair, because distant peaks are only expected to result from interactions with other molecules. Therefore, to build the feature vector, we used the cross-correlation profile that is cut off at different displacements, referred to as bin sizes (e.g. 2, 4, 10, and the length of the entire profile). The organism-specific final feature vector for a given metabolite-protein pair in SLIMP was obtained by concatenating the cross-correlation profiles over the different experiments (with selected bin sizes for displacement).

**Figure 2:**
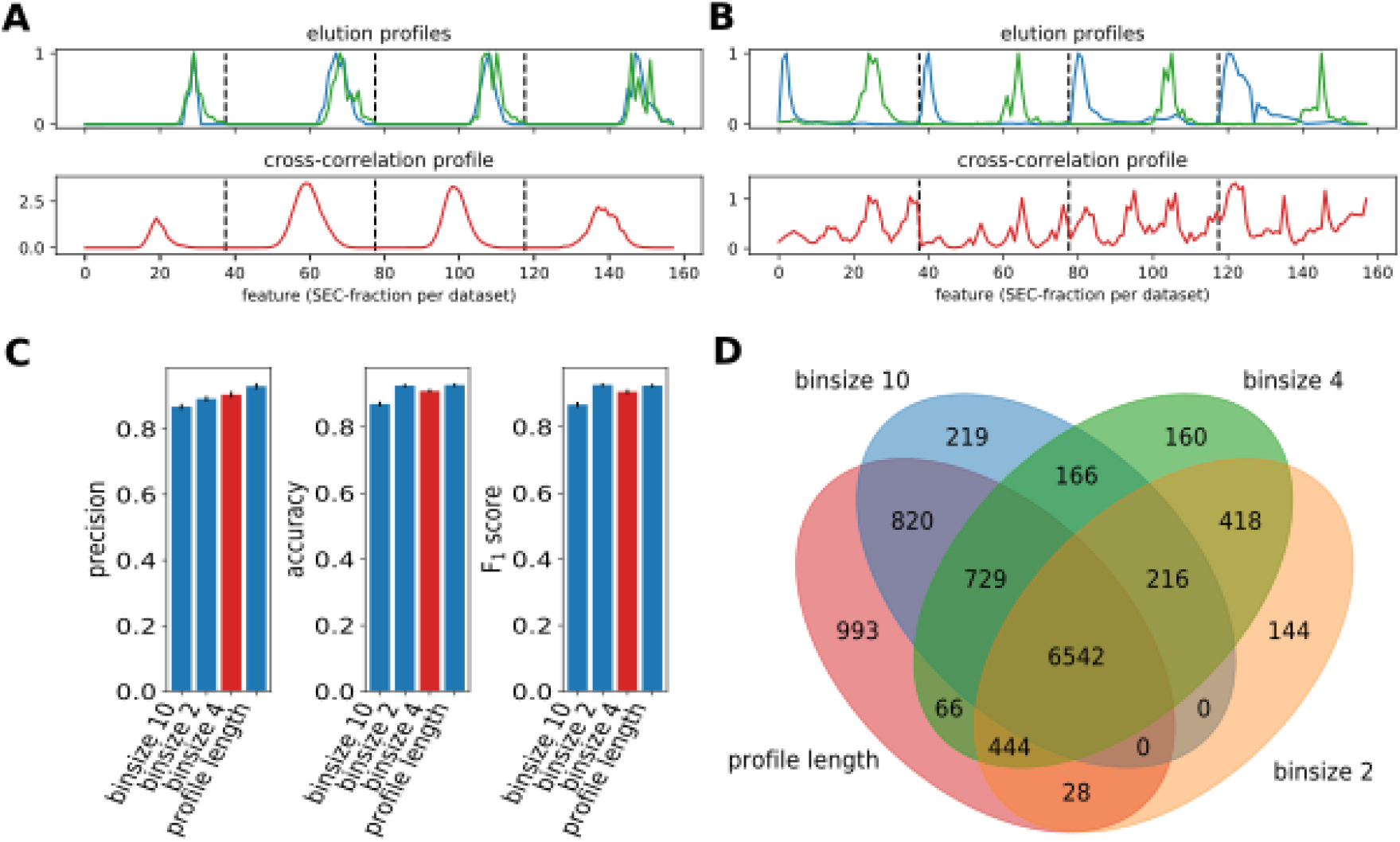
Feature engineering used in SLIMP. (**A**) Feature engineering for interacting molecules. Example of the feature engineering by calculating the full cross-correlation profile (red) from two given elution profiles of the co-eluting, interacting molecules malate dehydrogenase (green) and biotin (blue) (D. S. Szklarczyk 2016). (**B**) Feature engineering for not interacting molecules. Example of feature engineering for two not co-eluting, not interacting molecules Glyceraldehyde-3-phosphate dehydrogenase (green) and uridine (blue) with their resulting cross-correlation profile (red) (PubChem BioAssay AID: 2472). (**C**) Performance of the classifier on different bin sizes. Histograms show the values of various performance measures for the different bin sizes of the cross-correlation profiles. The selected bin size of 4 is marked in red. Means and standard deviations of the performance measures are calculated with 10-fold cross validation with 100 repetitions each. (**D**) Intersections of predicted interactions. Venn diagram showing the intersections of predicted interactions for different bin sizes on the full cross-correlation profile. All data from the performance on the *S. cerevisiae* experiment, profile pooling method “min”, linear support vector machine with regularization parameter C=0.1.

### Design of machine learning classifiers of metabolite-protein interactions

To train and evaluate the performance of a classifier using well-established machine learning approaches, the gold standard was split into training and test sets used in a 10-fold cross-validation. The classifiers varied with respect to the used pooling methods to combine experimental elution profiles over replicates of an experiment, bin sizes and normalization of the cross-correlation profile as well as the machine learning approaches employed.

To illustrate the effect of these factors on the performance of the classifiers, we first used linear support vector machines (SVM) with regularization on the features from *S. cerevisiae* for which the gold standard was of smaller size (Figure 2C,D). Most of the predicted interactions were shared when using cross-correlation profiles with different bin sizes. Predictions obtained when using cross-correlation profiles with displacement that corresponded to the length of the elution profile showed the worst agreement with every other tested bin size. This was likely a result of false positives due to over-weighting of larger displacements. In the prediction of MPIs we expect smaller displacements to exhibit better performance; however, a larger displacement may be favoured since the resulting classifier is expected to be more robust in a larger space of informative features. If a bin size includes larger displacements to peaks that are farther away, the classifier may suffer from over-fitting. In addition, a medium-sized bin size includes information about close elution peaks, excluding false positives caused by high cross-correlation due to broader, adjacent elution peaks which are included in too small bin sizes. Based on these arguments and since the performance of the classifier for different bin sizes was similar (Figure 2C), we decided to use a bin size of 4 in the remaining analyses.

Next, we investigated the effect of the different pooling methods (i.e. minimum, maximum, mean and median per fraction) used to combine the replicates of the elution profiles. Cross-validation results for the F_1_-score revealed that the performance of the linear SVMs with different pooling methods was similar (Figure 3A). The same could be concluded when other measures (i.e. precision, recall, accuracy, sensitivity and specificity) were used to assess performance, except for the higher standard deviation in sensitivity and specificity (Supplemental Figure S1). However, the predicted interactions vary: SVM classifiers trained on profiles pooled by taking the minimum resulted in the smallest, while that based on the mean yielded the largest number of predicted MPIs, (Figure 3B). Pooling profiles over replicates by the minimum of each fraction led to noise reduction, as peaks that were not observed in all replicates were excluded. Therefore, due to its robustness (and expected smaller number of false positives), we selected the minimum as a pooling method, at the cost of not identifying a larger number of interactions.

**Figure 3:**
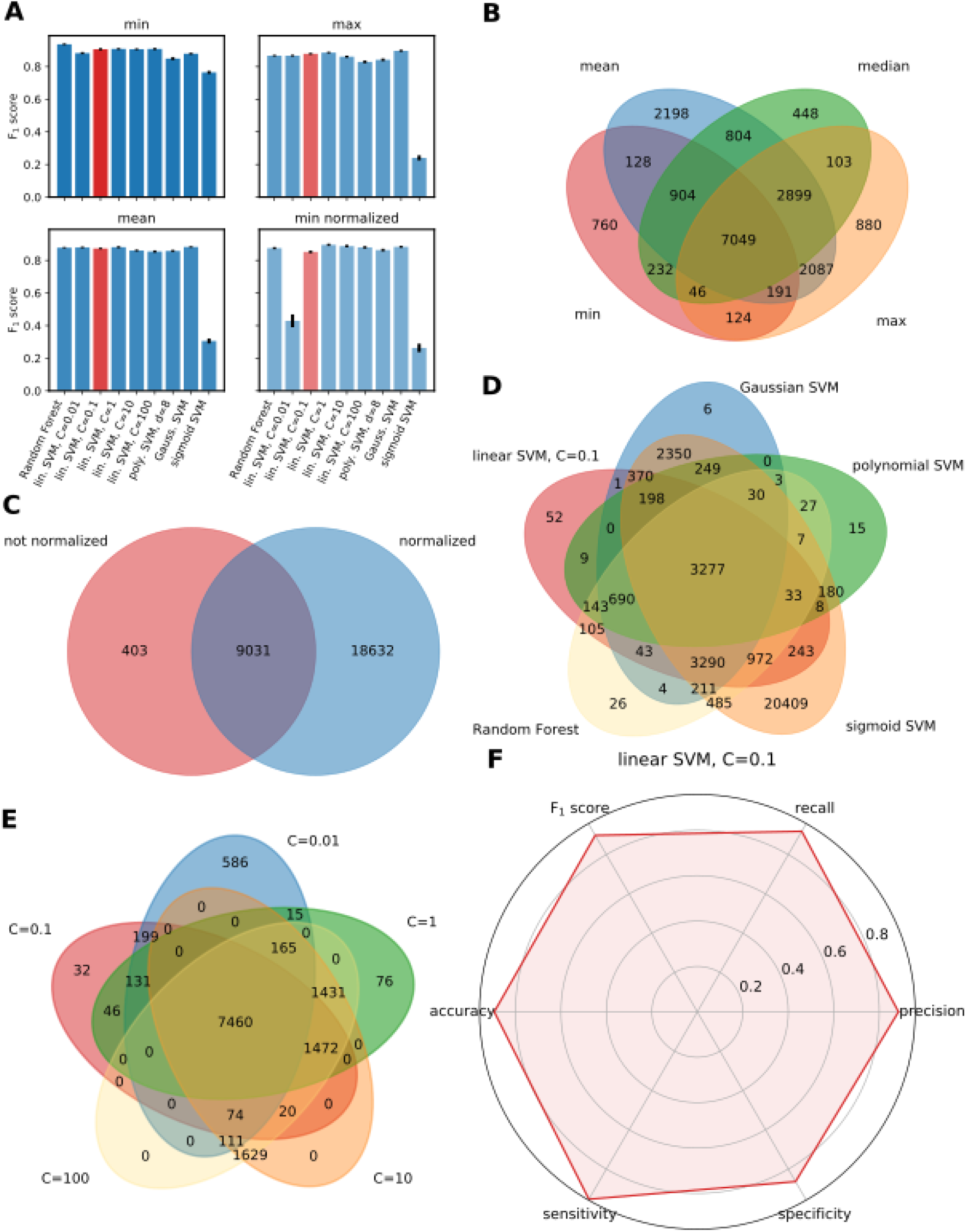
Effects of data- and algorithm-related factors on the performance of SLIMP. **(A)** Performance of different profile pooling methods. Histograms showing means and standard deviation of F_1_ scores for the performance with different profile pooling methods to combine repetitions of experiments (columns) for various classifiers (C=regularization parameter, d=degree of the polynomial kernel). (**B**) Venn diagram showing the intersections of interactions for different methods. (**C**) Intersection of interactions from classifiers trained on normalized vs. not normalized cross-correlation profiles to the maximum peak. (**D**) Comparison of the intersecting interactions for different classifiers. (**E**) Intersection of interactions for different regularization parameters C for the linear SVM. (**F**) Performance of the finally selected classifier. Radar chart displaying the value from 10-fold cross-validation of different performance measures for the selected classifier, SVM with linear kernel, regularization parameter C=0.1, pooling method minimum. Finally selected classifier is marked red. Abbreviations: lin.=linear, poly.=polynomial, Gauss.=Gaussian, norm.=normalized, SVM=support vector machine.

Further, we analysed the effect of normalization of the cross-correlation profiles to the maximum peak or the sum over the profile. We expect that normalization could help identify interactions between co-eluting molecules for which the intensity from mass spectrometry of one molecule type was much higher than the intensity of the other. However, this, too, might lead to false positives if the overlap is small. The comparison showed that most of the interactions predicted by the classifier without normalization were also predicted by the classifier based on normalized cross-correlation profiles (Figure 3C). As a result, we decided not to normalize the cross-correlation profiles, to reduce the number of false positive interactions.

Finally, we inspected the performance of the following machine learning approaches for classification: random forest, linear SVM with and without regularization or SVMs with polynomial, Gaussian radial basis function (RBF), and sigmoid kernel. In contrast to the other approaches, the SVM with a sigmoid kernel exhibited the largest number of predicted MPIs, resulting in a poor performance (Figure 3A,D). All the other approaches resulted in classifiers that performed equally well, reaching values of the F_1_-score greater than 0.8. Despite the best performance of the random forest we focussed on predictions of the SVM, since the random forest classifier tends to over-fitting as documented by the learning curves (Supplemental Figure S2). Further, given the size of the employed gold standard, we selected the linear SVM without any kernel as the classifier of choice. Regularization of the linear SVM had a small effect on prediction (Figure 3A,E). A smaller value for the regularization parameter C in our classification problem decreased the risk of over-fitting and resulted in a better performance according to the learning curve (Supplemental Figure S2, Methods).

Based on this evaluation, SLIMP is based on training a linear SVM with regularization using minimum-based pooled elution profiles without normalization of the cross-correlation profiles. Both classifiers showed excellent performance with respect to different performance measures on the data sets from *S. cerevisiae* (Figure 3F) and *A. thaliana* (Supplemental Figure S3).

### Representative value of the gold standard of metabolite-protein interactions

Apart from assessing different performance measures, there are other widely used characteristics to evaluate the performance of a classifier, including learning curves and receiver operating characteristic (ROC) curves. Learning curves provide insight on whether the gold standard (training data) is large enough and if the classifier suffers from under- or over-fitting (Pedregosa 2011). The small difference between the training and test scores implied that the SLIMP classifier did not suffer from over-fitting and does not exhibit high variance and high bias (Figure 4A). In addition, the learning curve implied that a bigger gold standard is not expected to increase the performance of our classifier estimator (Pedregosa 2011). We also investigated the diagnostic ability of our classifier by determining the ROC curve. The high value for the area under the ROC curve (AUC) indicated that the SLIMP classifier predicts well the presence and absence of interactions with low error rates (Figure 4B, upper panel).

**Figure 4:**
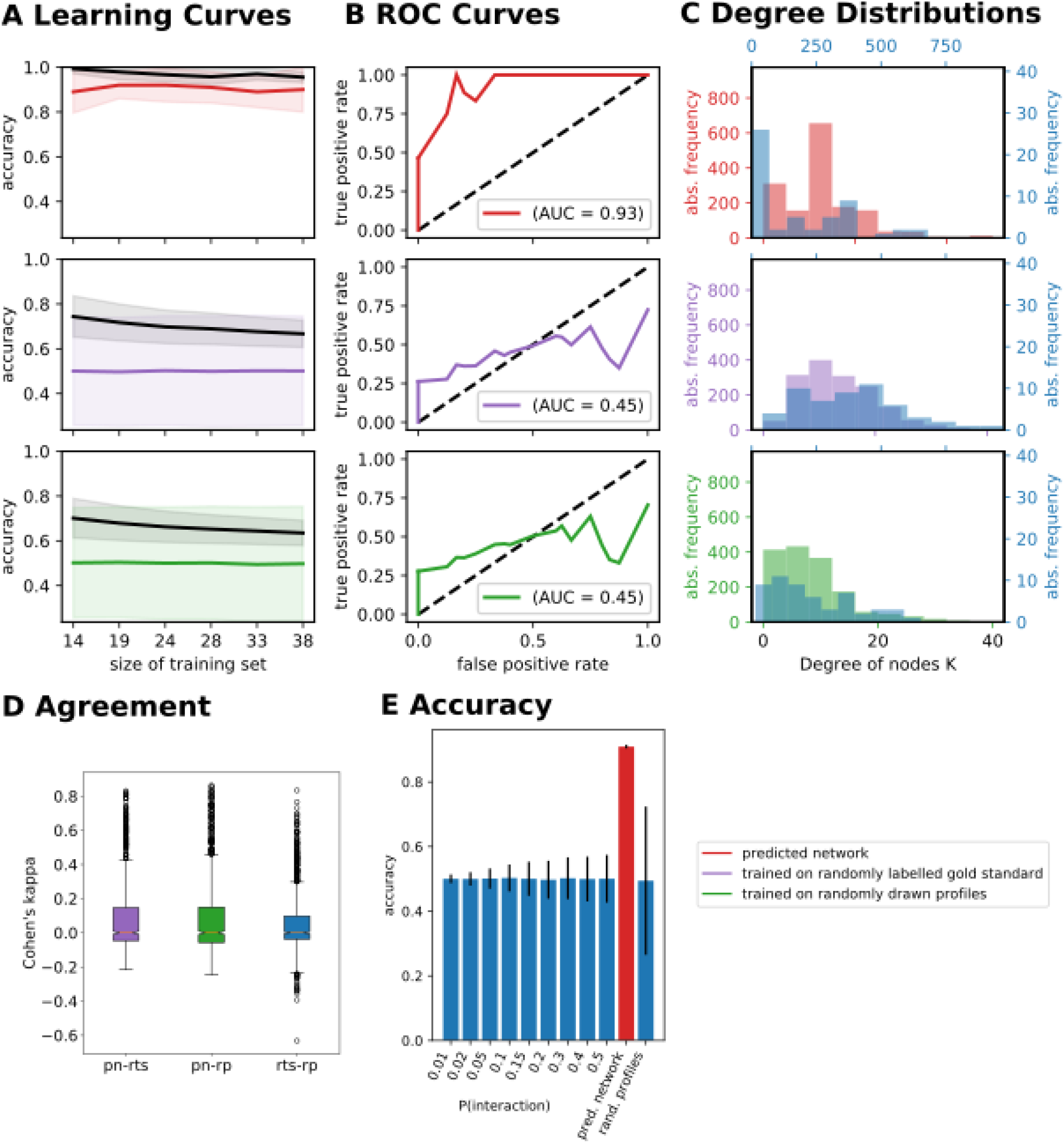
Comparison of predictions from classifiers trained on randomised data. (**A**) Learning curves for the classifier trained on the gold standard (red), a classifier trained with randomized labels on the training set (purple) and a classifier with randomly selected protein and metabolite profiles with random labels from the experiments (green, n=1000). Black lines represent mean accuracy on the training set and coloured lines on the test set, filled area shows the standard deviation from 10-fold cross-validation. (**B**) ROC curves for classifiers trained on the gold standard (red), the gold standard with randomized labels (purple) and randomly drawn profiles from the experiments (green, n=1000). Black dotted lines represent the diagonal. (**C**) Degree distributions (metabolites blue, proteins red for training on the gold standard, purple for training on the gold standard with randomized labels or green for training on randomly drawn profiles). (**D**) Boxplots display the agreement (Cohen’s kappa) of the predicted network (pn), network from randomly labelled gold standard (rts), and randomly drawn profiles (rp), n=1000.

Another test to investigate on the applicability of machine learning on the experimental data is to compare the performance of the SLIMP classifier to classifiers generated with randomized data. We decided to construct classifiers trained with the same gold standard, but with randomized labels (presence and absence of interactions, purple in Figure 4A–D). In addition, we trained a classifier on randomly drawn metabolite-protein pairs from the experimental data with random labels (green in Figure 4A–D). For every of the two types of random classifiers we took the average performance over 1000 classifiers, all trained with different randomized labels and profiles. The learning curves showed accuracy ~ 0.5, supporting random assignment of predicted interactions (Figure 4A). As seen in the high accuracy score on the training set, both classifiers trained on randomized data suffered from over-fitting. In addition, the poor performance is supported by the ROC curves, but not as a typical random assignment of the interactions (Figure 4B, dotted line). The shape of the ROC curves indicated that small false positive rates were associated with higher-than-random true positive rate; in contrast, larger false positive rates were associated with true positive rates that were smaller than those expected at random. As a result, the classifiers trained with random labels and profiles tended to predict absence of interactions resulting in a classification performance that is worse than random.

Further, we compared the prediction of the SLIMP classifier and the 1000 classifiers trained with randomized data. The results indicated no significant agreement between the predictions from the SLIMP classifier and those from a classifier trained on the gold standard with random labels (Figure 4D, purple) and randomly sampled profiles from all observed possible metabolite-protein pairs in the datasets (Figure 4D, green). Hence, the predictions made by the SLIMP classifier could not be obtained based on randomized data. We could also not see any agreement according to the Cohen’s kappa scores of the different random classifiers, as expected with completely random assignments (Figure 4D, blue), indicating that the profiles used in the gold standard match randomly drawn profiles from the pool of all observed MPIs. Summarizing the comparison to random classifiers, we showed that the gold standard of experimentally confirmed metabolite-protein (non)interactions are representative and that the engineered features contain valuable information about metabolite-protein interactions.

### Properties of the network of predicted metabolite-protein interactions

The previous findings corroborated the robustness of performance of the selected classifiers. Therefore, we next used the SLIMP classifiers along with 84,024 and 323,554 metabolite-protein pairs in *S. cerevisiae* and *A. thaliana* and predicted 9,434 and 20,148 MPIs, respectively. These MPIs can be represented as a bipartite network, with the two partitions composed of metabolites and proteins and edges denoting the MPIs. We first investigated the degree distributions of the metabolites and proteins as a seminal network property (Figure 4C, upper panel). We found that the degrees of metabolites in the network of predicted metabolite-protein interactions of *S. cerevisiae* did not follow a power-law distribution, in contrast to observations from analyses of networks assembled by mining of publically available databases of enzyme-metabolite inhibitory interactions (Alam 2017). On the other hand, the degrees of proteins tended to follow power-law, with the proteins that interact with the largest number of metabolites involved in folding and binding processes (e.g. chaperones and heat shock proteins). The degrees of proteins and metabolites in a network of predictions by classifiers trained on randomized data followed empirically a right-skewed distribution and covered a broader range of interactions than our predicted network (Figure 4C). Analogous findings were obtained for the network of predicted MPIs for *A. thaliana* (Supplemental Figure S4). Altogether, the analysis of the degree distributions of proteins and metabolites in the bipartite network of MPIs, predicted based on the selected SLIMP classifier, showed that this seminal network property differs from those of networks obtained from classifiers trained on randomized data, and may be considered biologically relevant.

### Validation and enrichment analysis of predicted metabolite-protein interactions

In the next step of our analysis, we focussed on identifying enriched interactions in the network of MPIs predicted by SLIMP. Different enzyme classes (according their EC numbers) showed different preferences to interact with metabolites of particular KEGG BRITE metabolite classes (Figure 5A). For instance, all enzymes have significantly enriched interactions to coenzymes. On the other hand, proteins with non-enzymatic function, many of which are involved in DNA-binding, showed enriched interactions to nucleotides and nucleosides. As in the network displayed in Figure 5A, some well-known interactions were supported by our predictions when we inspected the enrichment of interactions between protein and metabolite KEGG BRITE classes (Figure 5B).

**Figure 5:**
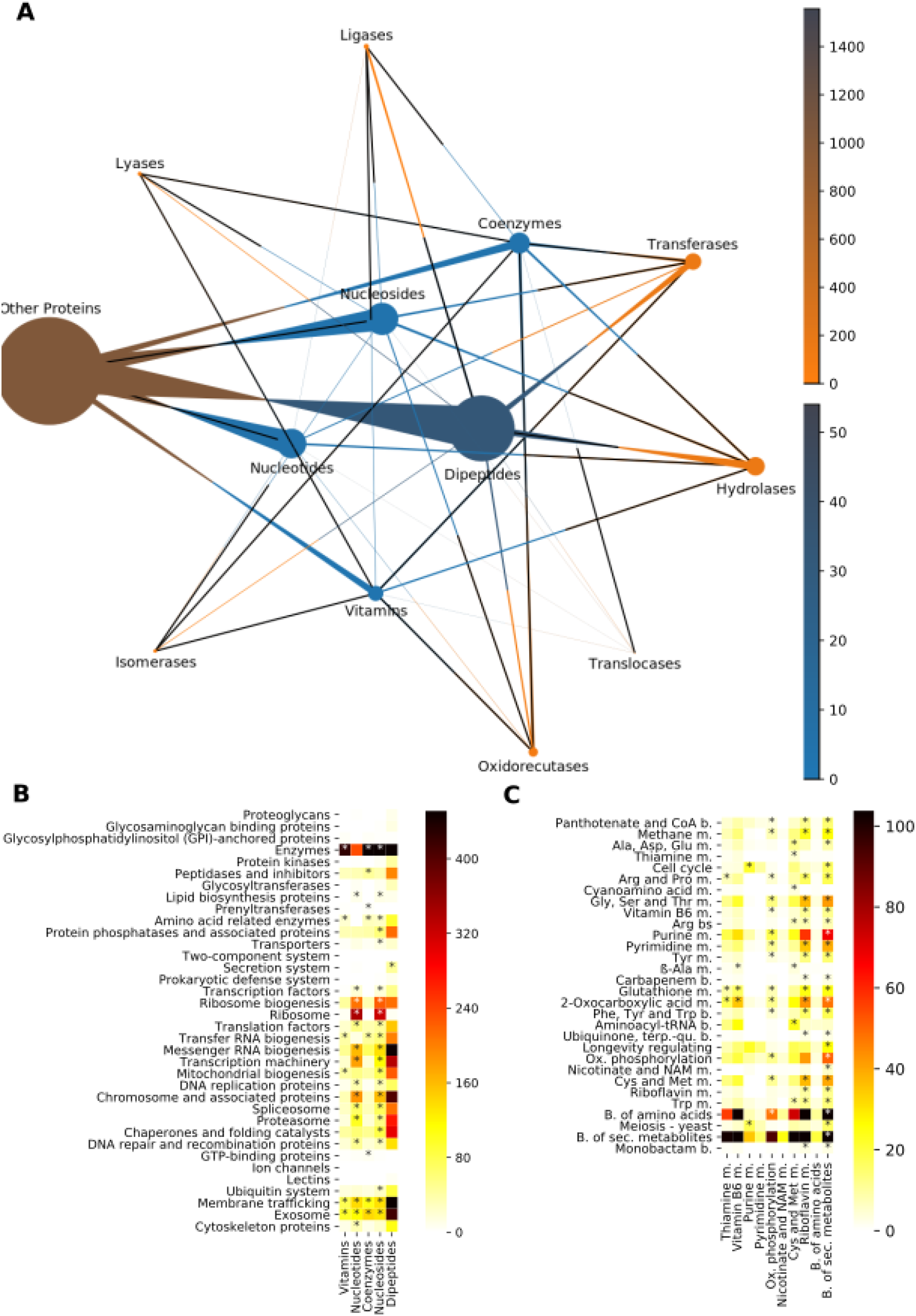
Enrichment analysis of the metabolite-protein interactions predicted from SLIMP. (**A**) Bipartite network of MPIs between EC-classes and KEGG metabolite classes. The size of a node describes the fraction of all determined interactions, the colour describes the number of molecules in the given class and the triangular directed edges how many of the interactions from the given source class are with the target class. Black edges symbolize significantly larger tendency of interactions from source to target class than expected by chance, α=0.05. (**B**) Heatmap for number of interactions between proteins of different BRITE classes (rows) and KEGG metabolite classes (columns). (**C**) Heatmap for number of interactions from proteins (rows) to metabolites (columns) between different pathways, m.=metabolism, b.=biosynthesis. Asterisks mark significantly more interactions than expected by chance, α=0.05, p-values corrected for multiple testing using Bonferroni-correction.

**Figure 6:**
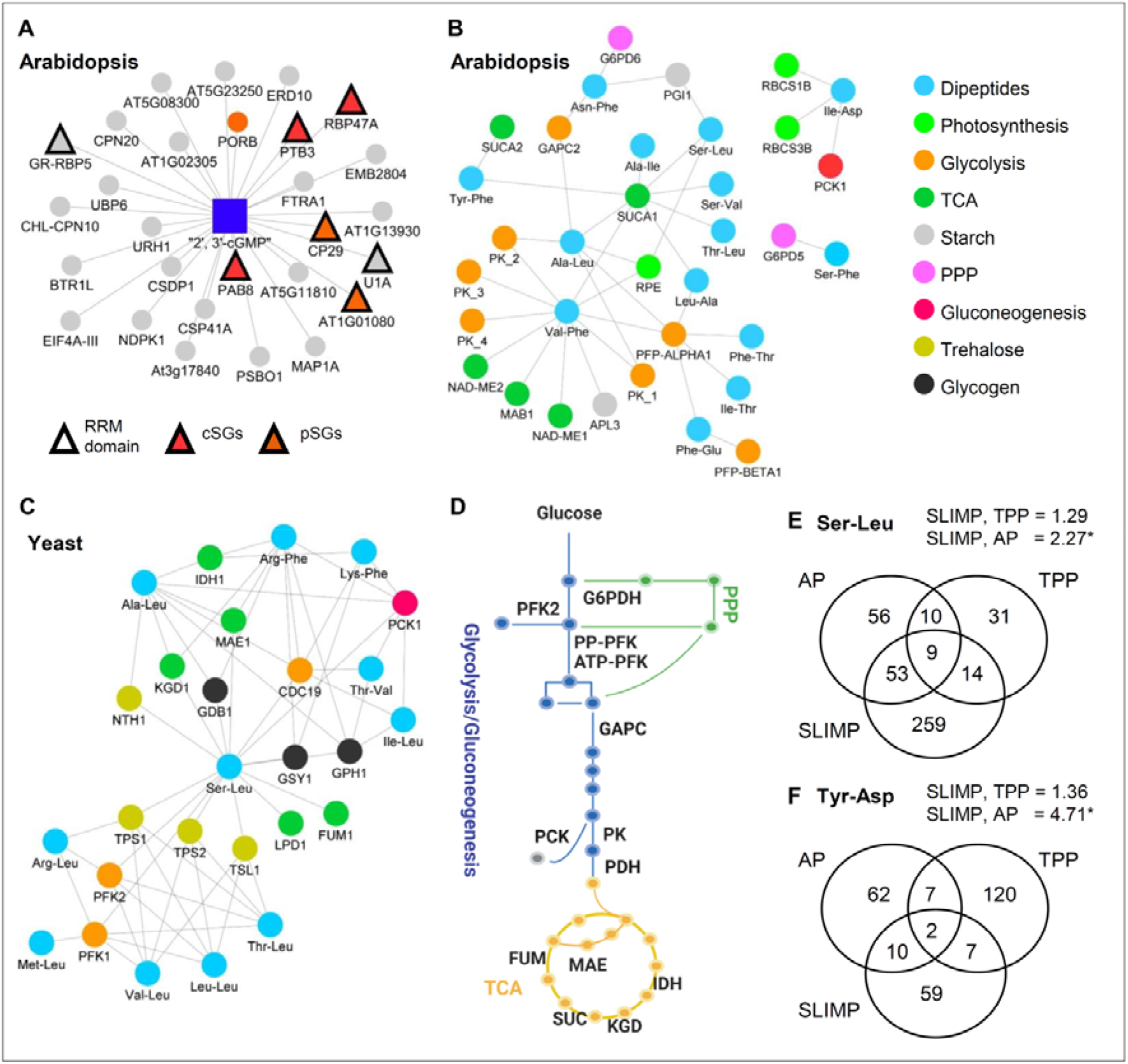
Novel interactions between 2’, 3’-cNMP, dipeptides and proteins revealed by SLIMP. (**A**) 31 high-confidence 2’, 3’-cGMP targets delineated by SLIMP. RRM, RNA binding motif; cSG, cytsolic stress granule, pSG, plastidial stress granule. (**B, C**) Sub-network depicting interactions between dipeptides and enzymes from the central carbon metabolism in (B) Arabidopsis and (C) yeast. (**D**) Schematic representation of the main pathways in the central carbon metabolism (glycolysis, gluconeogenesis, TCA cycle and pentose phosphate pathway). Enzymes found among dipeptides’ high confidence protein interactors in either Arabidopsis or yeast are highlighted. (**E, F**) Comparison of targets delineated by SLIMP, affinity purification (AP) and thermal proteome profiling (TPP) for (E) Tyr-Asp in arabidopsis (all interactions predicted by SLIMP) and (F) Ser-Leu in yeast (SLIMP prediction probability for interaction > 80 %). (A-C) were constructed using Cytoscape, (D) was prepared using Biorender.com.

For instance, our predictions indicated that enzymes tend to interact with coenzymes and their precursors, proteins involved in transcription and translation preferably interact with nucleosides. Nucleotides were found in more interactions with proteins from synthesizing processes (Figure 5C). Interestingly, we also found that most interactions are found across and not within pathways. For *A. thaliana* similar tendencies were observed; for instance, enzymes tend to interact with coenzymes and their precursors, while translation factors favour interactions with nucleosides; moreover, many MPIs tend to bridge different pathways (Supplemental Figure S5). These results represent well-known characteristics of MPIs, which support the accuracy and value of the interactomes predicted by SLIMP.

### Novel targets of 2’, 3’ cyclic nucleotides and dipeptides identified by SLIMP

Following the enrichment analysis, we focused on the two groups of metabolites present in the SLIMP dataset, with recently described regulatory functions, namely: 2’, 3’ cyclic nucleotides (2’, 3’-cNMPs) and proteogenic dipeptides. 2’, 3’-cNMPs are evolutionarily conserved RNA degradation products, which accumulate under stress conditions (Jackson 2015). It has been shown that 2’, 3’-cAMP, and 2’, 3’-cGMP bind to the RNA Recognition Motif (RRM) of the Arabidopsis Rbp47 protein (M. L.-B. Kosmacz 2018). RRM domain is conserved among the eukaryotes, and proteins such as Rbp47 characterized by RRM motif and prion-like unstructured domain are known to play a central role in the stress granules (SGs) assembly (Protter 2016). In agreement with the binding data, 2’, 3’-cAMP treatment-induced cytosolic SGs assembly in the Arabidopsis seedlings (M. L.-B. Kosmacz 2018). Corroborating the published work, seven of the 31 high-confidence 2’, 3’-cGMP targets predicted by SLIMP (prediction probability for interaction > 80 %) are characterized by RRM domain, Rbp47 among them, which is 12.8-times more than expected by chance. The list also contains a plastidial protein CP29, speculated to play a similar role to the Rbp47 for the assembly of plastidial SGs (Chodasiewicz 2020). Furthermore, of the 31 proteins, six were shown to sequester in either cytosolic or plastidial SGs (Chodasiewicz 2020, M. G. Kosmacz 2019). Based on the SLIMP results, we now hypothesize that 2’, 3’-cNMP facilitate both cytosolic and plastidial SGs formation *via* binding to the RRM motif of the SGs proteins.

Similar to 2’, 3’-cNMPs, dipeptides are degradation products and accumulate under conditions associated with proteolysis, such as heat stress in plants (Thirumalaikumar 2021) and glucose depletion in yeast (M. V. Luzarowski 2021). We have demonstrated that dipeptides bind and regulate the activity of the glycolytic/gluconeogenic enzymes, and supporting the binding data, dipeptide feeding leads to acute metabolic changes. SLIMP predicted dipeptide-protein interaction network (prediction probability for interaction > 80 %) comprising of 607 interactions between 31 dipeptides and 221 proteins in *A. thaliana* and 1951 interactions between 14 dipeptides and 516 proteins in *S. cerevisiae.* Based on the published regulatory interactions between dipeptides and central metabolism, we focused specifically on interactions between dipeptides and enzymes in carbon metabolism. The predictions of the dipeptide-enzyme interaction network by SLIMP based on data from Arabidopsis comprised glycolytic enzyme glyceraldehyde-3-phosphate dehydrogenase (GAPDH/GAPC) and gluconeogenic enzyme phosphoenolpyruvate carboxykinase (PCK1) (Moreno 2021) We have already demonstrated that Tyr-Asp inhibition of the GAPC activity is associated with the diverting of glycolytic flux towards pentose phosphate pathway (PPP) and NADPH production(Moreno 2021). Interestingly, among the new, putative targets identified by SLIMP we found Glucose-6-phosphate dehydrogenase (G6PDH) whose regulation may further contribute to the observed metabolic phenotype, since the concomitant regulation of the GAPDH and G6PDH activities controls the flux into PPP in bacteria, yeast, and human cell lines (Christodoulou 2019). Additional enzymes identified in our analysis in both Arabidopsis and yeast included phosphofructokinase, pyruvate kinase, and NAD-dependent malic enzyme, all of which are focal points for metabolic regulation. Based on the SLIMP results, we now hypothesize a multisite regulation of carbon metabolism by dipeptides.

Finally, we compared the MPIs predicted by SLIMP with interactions retrieved previously using affinity purification (AP) and thermal proteome profiling (TPP) for dipeptide Tyr-Asp in Arabidopsis and dipeptide Ser -Leu in yeast. In both cases, we found a significant overlap between SLIMP and AP targets lists (Fisher exact test, *p*-value <0.05) but not between SLIMP and TPP, which may reflect the common, chromatographic principle behind SLIMP and AP. In contrast, TPP looks for the difference in a protein melting temperature upon ligand binding, and will miss interactions that elicit no or only weak effect on a protein thermal stability. However, the interactors found in at least two of the three approaches constitute priority for future characterisation, because of the inherit differences between the three methods interactors identified by one approach, only, should not be dismissed.

In summary, SLIMP analysis retrieved known interactions, validating our approach, but also generated novel hypothesis to be tested experimentally.

## Conclusion

Here we designed, evaluated, and tested the performance of a supervised approach for prediction of metabolite-protein interactions termed SLIMP. Our approach relies on combining SEC and LC-MS to obtain metabolomic and proteomic profiles, and can be applied on organisms of different complexity, as shown for *S. cerevisiae* and *A. thaliana*. With the advances in mass spectrometry and development of techniques for identification of metabolite-protein interactions, SLIMP provides a scalable strategy to predict the entirety of metabolite-protein interactions. Key to this prospect is the availability of high-quality gold standard, particularly of negative instances which are often missing or are not reported in biological studies. While SLIMP predicts metabolite-protein interactions with good quality, as assessed by different performance measures, it does not distinguish between interactions of a protein in monomeric state and those with protein complexes. Nevertheless, our data sets and approach can be adapted to predict protein-protein interactions. In addition, SLIMP does not categorize the type of interactions (e.g. activating, inhibiting, structural, and non-functional). Provided detailed gold standard, future efforts will be directed at using SLIMP in combination with metabolic networks to allow for more refined predictions about the putative function of MPIs.

The comprehensive comparative analyses with state-of-the-art approaches for identification of metabolite-protein interactions indicated significant agreement of predictions from SLIMP with findings from an independent experimental procedure. Further, we corroborated the quality of the network of MPIs predicted from SLIMP with enrichment and structural analyses as well as cross-species comparisons of new MPIs involving dipeptides. Therefore, since there exists no standard approach to obtain a global map of MPIs *in vivo*, SLIMP can be viewed as a valuable approach for generation of hypotheses about such interactions.

## Materials and Methods

### Method Details

The data acquisition is based on size exclusion chromatography and further proteomic and metabolomic analyses, as in the PROMIS approach (Sokolowska 2019, D. S. Veyel 2018, M. V. Luzarowski 2021). In contrast to PROMIS, SLIMP is a supervised approach which relies on carefully engineered features that combine the SEC elution profiles. The overall procedure was applied to different cell-lysates of *A. thaliana* and *S. cerevisiae*.

### Plant and yeast material grow conditions

*Arabidopsis thaliana Ler* cell suspension culture, was grown in MSMO (Murashige and Skoog basal salts with minimal organics, Sigma) medium, in a in constant light (c. 80 μmol m–2 s–1), at 21 °C, on orbital shaker (110–120 rpm). Cells were harvested at the logarithmic growth phase (7 days after last passage) by filtration and immediately frozen in liquid nitrogen. As described in (Veyel et al., 2018). *Arabidopsis thaliana* Col-0 seedlings (1.5 mg seeds, c. 100 plants) were grown in sterile liquid cultures (100-ml Erlenmeyer glass flasks) in 35 ml ½ MS (half strength, Murashige and Skoog, Sigma) liquid medium with 1% sucrose. The seedlings were kept shaking for 10 days on orbital shakers (110–120 rpm), in constant light (c. 80 μmol m–2 s–1) and temperature (21°C) conditions. Plants were harvest and snap frozen in liquid nitrogen. *Arabidopsis thaliana* Col-0 rosettes were grown on soli in a controlled condition, in a growth chamber (Phytotron GmbH) with 12-h light at 120 μmol m-2 s-1, in the constant temperature of 21°C. Four weeks old plants were harvest and snap frozen in liquid nitrogen. The *Saccharomyces cerevisiae* strain YSBN2 was cultivated at 30 °C with moderate shaking (120-140 RPM) using Innova Shakers. Cultures were started using a single colony grown on YPD (Yeast extract Peptone Dextrose, Sigma) plate. After 6 (logarithmic phase), 24 (diauxic phase) and 72 (stationary phase) hours of cultivation, cells were collected by centrifugation (4,000 g, 4 °C, 20 min), washed with AmBIC buffer (50 mM ammonium bicarbonate, 150 mM NaCl, 1.5 mM MgCl_2_), transferred to 50 mL tube and centrifuged again (4,000 g, 4 °C, 20 min). The yeast pellet was then snap frozen in liquid nitrogen.

### Experimental generation of elution profiles

PROMIS experiments was performed as described previously in (D. S. Veyel 2018) (M. V. Luzarowski 2021). Briefly, native lysate containing endogenous protein- protein and protein-metabolites complexes corresponding to 40 mg of protein, was separated by size-exclusion chromatography (SEC) with Sepax SRT SEC-300 21.2 × 300 mm column (Sepax Technologies, Inc., Delaware Technology Park, separation range 1.2 mDa to 10 kDa) with the ÄKTA explorer 10 (GE Healthcare Life Science, Little Chalfont, UK). 36-40 1-mL protein containing fractions covering sizes from 13 to 6037.7 kDa were collected, snap frozen in liquid nitrogen and lyophilized.

#### Liquid-Chromatography Mass Spectrometry

Metabolites and Proteins from the lyophilized fractions were extracted using a methyl-tert-butyl ether (MTBE)/methanol/water method (Giavalisco 2011), where, molecules are simultaneously separated into organic phase (lipids), aqueous phase (polar and semi-polar metabolites) and protein pellets. Aqueous phase and protein pellets were dried in a SpeedVac and subjected metabolomic and proteomic analysis as described by Sokolowska, 2019. Briefly, the dried aqueous phase was suspended in 100 μL water. Samples were analysed by ACQUITY UPLC I Class (Waters) coupled with Exactive mass spectrometer (Thermo Fisher Scientific) in positive and negative ionization modes. The mobile phases consisted of 0.1% formic acid in water (Solvent A) and 0.1% formic acid in acetonitrile (Solvent B) and the gradient ramped as fallows: 1 min 1% B, 11 min 1% to 40% B, 13 min 40% to 70% buffer B, then 15 min 70% to 99% B, followed 2 min washout with 99% B. Mass spectra were acquired using following settings: mass range from 100 to 1500 m/z, resolution set to 25,000, loading time restricted to 100 ms, AGC target set to 1e6, capillary voltage to 3 kV with a sheath gas flow and auxiliary gas value of 60 and 20, respectively. The capillary temperature was set to 250 °C and skimmer voltage to 25 V. Acquired chromatograms were further processed with Expressionist Refiner MS (http://www.genedata.com) and obtained metabolite clusters were matched to in-house reference compounds library for annotation. Protein pellets obtained after MTBE extraction were resuspended in 50 μL denaturation buffer (6 M urea, 2 M thiourea in 40 mM ammonium bicarbonate). Reduction of cysteines, alkylation and enzymatic digestion using LysC/Trypsin Mix (Promega Corp., Fitchburg, WI) followed by desalting of a digested peptide was performed according to the protocol described in 1. Dried peptides were resuspended in MS loading buffer (3% ACN, 0.1 % FA) and measured with Q Exactive HF (Thermo Fisher Scientific) coupled to a reverse-phase nano liquid chromatography ACQUITY UPLC M-Class system (Waters). Equivalent of 1ug of proteins was injected per run and the gradient ramped from 3.2% ACN to 7.2% ACN over 20 min, then to 24.8% ACN over next 70 min and to 35.2% ACN over next 30 min, followed by a 5 min washout with 76% ACN. The MS was run using a data dependent acquisition method. Full scans were acquired at a 120,000 resolution, m/z ranging from 300.0 to 1600.0, a maximum fill time of 50□ms and an AGC target value of 3e6 ions. Each dd-MS2 scan was recorded at the resolution of 15,000 with an AGC target of 1e5, maximum injection time 100 ms, isolation window 1.2 m/z, normalized collision energy 27 and the dynamic exclusion of 30 sec. Raw proteomics files were analyzed using MaxQuant software with Andromeda an integrated peptide search engine with a fallowing setting: maximum of two missed cleavages were allowed and threshold for peptide validation was set to 0.01 using a decoy database, methionine oxidation and N-terminal acetylation was considered as variable modification while cysteine carbamidomethylation as a fixed modification. The minimum length of peptide was set to at least seven amino acids. Moreover, following options were selected: “label-free quantification” and “match between runs”. Peptides were identified for Arabidopsis data sets using: *A. thaliana* Uniprot protein sequences (UP000006548) or Arabidopsis TAIR database (Version 10, The Arabidopsis Information Resource, www.Arabidopsis.org) and *Saccharomyces cerevisiae* protein database, Uniprot (UP000002311).

### Feature Engineering for Machine Learning

Further analysis aimed to determine whether two profiles, one from a protein and the other from a metabolite, show a co-elution behaviour. The relation between molecules (interaction or no interaction) was predicted by testing every protein against every metabolite on co-elution.

#### Profile Normalization

Comparability of profiles with MS intensity from different molecules was achieved by normalization of every profile to its maximum intensity. Normalization to the maximum intensity was chosen over normalization to the sum of the profile to address the challenge resulting from protein-protein interactions. More specifically, it is not only that metabolite-protein complexes elute through the SEC column, but it also contains protein-protein-complexes. If a protein multimerizes or assembles with other proteins, and therefore shows several peaks in the elution profile, normalization to the sum of the profile would decrease normalized peak intensities compared to monomeric proteins. With normalization to the maximum of each profile the peak intensity becomes independent of the total number of peaks per profile

#### Profile Pooling

For all the experiments with the different cell lysates replicates were made to provide reproducibility and the profiles of every molecule were combined over replicates using a pooling method (one of mean, median, minimum, or maximum). The pooling method of choice was selected *a posteriori* based on the by the performance of the classifier.

#### Data Analysis Experiments

To increase the robustness of made predictions we concatenated profiles of each molecule from different datasets. We designed two data analysis experiments with the datasets from different experiments. One experiment was performed on metabolite-protein pairs in the three datasets from *A. thaliana*, the second on all four datasets from *S. cerevisiae*.

#### Combination of Protein and Metabolite Profiles

For the application of machine learning approaches the profiles of every metabolite-protein pair were combined. Theoretical considerations led to the use of the cross-correlation profile of two molecules.

Cross-correlation is a measure of similarity, which gives a function of the displacement of one series relative by another (Bracewell 1986). For finite, discrete series *p* and *m* describing the elution profiles of proteins and metabolites, respectively, the cross-correlation function (*p* ⋆ *m*), giving the displacement of the protein profile by the metabolite profile is defined as following (C. Wang 2019):

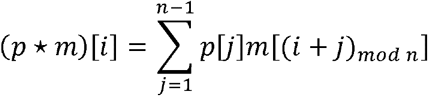

where *mod* defines the modular arithmetic operation and [*x*] accesses the *x^th^* element of the series, in our case the normalized MS-intensity of an elution profile.

With no displacement, two profiles from co-eluting molecules should show highest similarity (overlapping elution peaks), whereas further displacement should reduce similarity. Molecules that do not co-elute, show overlap of their elution peaks only after profile displacement and exhibit low or no similarity without displacement (Figure 2). We assume that larger displacements do not hold information about interactions, since more distant peaks may induce false interpretations due to interactions with other molecules and therefore tested different maximum displacements, further referred to as binsizes (4,10, and total profile length). The resulting cross-correlation profiles were concatenated over the included experimental datasets per data analysis experiment.

The shape of the cross-correlation profile denoting an interaction between two molecules is expected to have one peak at zero displacement per dataset. However, we cannot estimate the mathematical parameters describing the peaks, so we used machine learning classifiers which are able to mimic the shape and assign values to the unknown parameters.

The cross-correlation profiles served as feature vectors for these different machine learning classifiers. Every element of the vector describing the cross-correlation profile corresponds to the sum of fraction-wise products of displaced protein and metabolite elution profiles. The cross-correlation profiles cut off at the different bin sizes were used as engineered features for the classifiers and the best performing classifier was selected.

### Prediction of Interactions Using Machine Learning

The different classifiers learned with the engineered feature vectors from given metabolite-protein pairs present in the gold standard with their respective relation (presence or absence of interaction) to identify patterns of feature vectors for interactions and non-interactions. A classifier trained with the gold standard is able to make predictions about interactions on metabolite-protein pairs with unkown relations.

Based on the amount of available training data (Supplemental Table S3), we compared different support vector machines (SVM) and random forest classifiers. These binary classifiers were trained with cross-correlation profiles of metabolite-protein pairs with known relation extracted from databases and estimated a decision function to predict the presence or absence of interactions by categorizing metabolite-protein pairs into “interacting” or “not-interacting” per our experimental data in combination with published evidence.

#### Support Vector Machines

A support vector machine constructs a multidimensional space with as many dimensions as given features, in our case elements in the cross-correlation profile. Every cross-correlation profile is represented in this space as a point with its profile elements as coordinates and the SVM finds a separating hyperplane with maximized margin between the agglomerations of these points from interacting and not interacting profiles. The hyperplane assigns weights to every feature. Features describing no or low displacement between a metabolite and protein profile obtained positive weights, features describing medium displacements were characterized by negative weights. The decision function describing the hyperplane is calculated by choosing a function that minimizes the training error on the gold standard. Regularization in the SVM chooses a function that additionally minimizes the norm of the function to reduce the sum over all feature weights and avoid overfitting on the training data (Bishop 2006). A higher value of the regularization parameter C makes the classifier more sensitive to outliers and therefore prone to over-fitting, whereas a lower value forces the classifier to be more generic and prone to under-fitting. We also tested the effect of various kernel functions that extend the multidimensional space by an additional dimension described via a similarity function. To this end, we used Gaussian radial basis function, sigmoid kernel and polynomial with maximum degree d=8 such that the kernel function has as many peaks as datasets are aligned to match the shape of the estimated cross-correlation profile (Figure 2A). A SVM is well applicable to our cause with little training data and high-dimensional feature space (Pedregosa 2011).

#### Random Forests

Another classifier we used is the random forest classifier, which builds many decision trees that each decide on selected features from the cross-correlation profile over presence or absence of interactions (Breiman 2001). The overall categorization of the metabolite-protein relation is done by a majority vote over all decision trees. Random forest classifiers provide weights to the features used and are therefore theoretically suitable to make predictions from our data.

#### Creation of the Gold Standard

The machine learning classifiers need to be trained based on features for the instances in the gold standard to predict presence or absence of MPIs. The classifiers trained with the cross-correlation profiles assigned labels which state whether the two molecules do or do not interact. To obtain balanced classifiers that show no bias towards any tendency of relation (interaction or no interaction), the training set is assembled of a negative and a positive set of equal size, containing not interacting and interacting metabolite-protein pairs, respectively.

#### Positive Set

The positive set was extracted from the STITCH database (D. S. Szklarczyk 2016) and PubChem Bioassays (Y. S. Wang 2013). The database assigns scores between 0 and 1000 to interactions between proteins and small molecules which state the confidence of the interaction. For the construction of the positive set only interactions with a score of 800 (80 % confidence) or above were included. Additionally, PubChem Bioassays was used for the acquisition of the positive set to access binding assay data for orthologous proteins in *Homo sapiens*, *Escherichia coli* and *Saccharomyces cerevisiae* and interacting metabolites. Since these databases consist of mostly experimental *in vitro* data, which may differ from *in vivo*- like conditions, we manually selected profiles which respect the assumption of co-elution between molecules to create a gold standard.

#### Negative Set

The negative set, describing absence of interactions is as important as the positive set for balanced training data. Negative results are often neglected in research and their value is underestimated, although they are highly valuable for supervised machine learning. However, it is still difficult to find research publishing absence of interactions. The PubChem Bioassays database keeps track of binding assays asserting absence of interactions, and not interacting metabolite-protein pairs were extracted via orthologous proteins in *Homo sapiens*, *Escherichia coli* and *Saccharomyces cerevisiae*.

For the search in PubChem Bioassays proteins were annotated to orthologous groups using the STRING database. This database uses hierarchical orthology imported from the eggNOG database (Huerta-Cepas 2016) to collect similar proteins between and within species (D. M. Szklarczyk 2016). An orthologous group was assumed to interact with a metabolite only, if all proteins with interaction records in the database of that orthologous group were interacting with the metabolite.

### Quantification and Statistical Analysis

#### Data sets employed in prediction of metabolite-protein interactions

The provided datasets (Supplemental Datasets S1-S2) only include all intersecting proteins and metabolites.

#### Design of machine learning classifiers of metabolite-protein interactions

The mean values and standard deviation for different performance measures (precision, recall, F_1_-score, accuracy, sensitivity, specificity) on all used classifiers are collected in the Supplemental Table SourceDataForFig3AF.

#### Representative value of the gold standard of metabolite-protein interactions

Learning and ROC curves were calculated for using 10-fold cross-validation, for classifiers trained on randomly selected profiles on 1000 repetitions. The obtained mean and standard deviation values for the SLIMP classifier can be found in Supplemental Table SourceDataForFig4. For the calculation of learning curves 80% of the gold standard were used for training, whereas the remaining 20% were reserved for the test set.

#### Properties of the network of predicted metabolite-protein interactions

Detailed values for the agreement between predictions of the SLIMP classifier and classifiers based on randomized training data as well as accuracies are summarized in the Supplemental Table SourceDataForFig4. The code for the generation of randomized networks is provided (Data Availability).

#### Validation and enrichment analysis of predicted metabolite-protein interactions

The network of interactions between metabolite and protein classes was constructed for metabolites with annotated KEGG metabolite classes and proteins with annotated EC numbers (Figure 5A). Proteins without annotated EC numbers were classified as ‘Other Proteins’. In the network the size of every node represents the number of interactions the given class is involved and the colour the number of annotated molecules in the class. The width of the triangular edges represents how many interactions of a given class are present towards another class. If significantly more interactions from one class to another were found, the centre of the edge was marked black. The significance was determined using Fisher’s exact test with Bonferroni correction for multiple testing on a significance level of 0.05.

We assigned BRITE classes and KEGG pathways to metabolites and proteins to construct heatmaps of interaction tendencies among classes and pathways (Figure 5B,C). (Macro-) molecules without annotated class/pathway were excluded from the analysis. Interactions between classes that are found significantly enriched are marked with a star. To calculate the significance we used Fisher’s exact test with Bonferroni correction for multiple testing on a significance level of 0.05. The number of interactions and non-interactions among classes and pathways used for the generation of heatmaps are summarized in Supplemental Table SourceDataForFig5.

## Author Contributions

ZN and AS designed the research. BZ designed, performed all computational analyses and wrote the manuscript. ZN designed the computational analyses and wrote the paper. AS designed PROMIS separations, analyzed metabolite data and wrote the manuscript. ML performed PROMIS in *Saccharomyces cerevisiae*, analyzed protein and metabolite data, and edited the manuscript. ES performed PROMIS experiment from *Arabidopsis thaliana* rosettes, analyzed protein and metabolite data from Arabidopsis PROMIS experiments, and edited the manuscript. DS, analyzed protein and metabolite data, and edited the manuscript. EL preformed PROMIS experiment from *Arabidopsis thaliana* seedlings under MK supervision.

## Data Availability

### Lead Contact

Further information and requests for resources and reagents should be directed to and will be fulfilled by the Lead Contact, Zoran Nikoloski

### Materials Availability

No new unique reagents were generated in this study.

### Data and Code Availability

- Computer scripts: https://github.com/bobbinf/slimp (approximated runtime: 25 h for creation of gold standard and prediction of interactions for S. cerevisiae on Linux kernel version 5.5.16-200.fc31.x86_64, operating system GNU/Linux, 8 Intel(R) Core(TM) i7-3720QM CPU @ 2.60 GHz and 16GB RAM

## References

Abdi, H. “RV coefficient and congruence coefficient.” Encyclopedia of measurement and statistics, 2007: 849–853.

Alam, M. T., Olin-Sandoval, V., Stincone, A., Keller, M. A., Zelezniak, A., Luisi, B. F., Ralser, M. “The self-inhibitory nature of metabolic networks and its alleviation through compartmentalization.” Nature communications, 2017: 1–13.

Aoki-Kinoshita, K. F., Kanehisa, M. “Gene annotation and pathway mapping in KEGG.” Comparative Genomics, 2007: 71–91.

Bishop, C. M. Pattern recognition and machine learning. Springer, 2006.

Bracewell, R. N., Bracewell, R. N. The Fourier transform and its applications (Vol. 31999). New York: McGraw-Hill, 1986.

Breiman, L. “Random forests.” Machine learning, 2001: 5–32.

Chodasiewicz, M., Sokolowska, E. M., Nelson-Dittrich, A. C., Masiuk, A., Beltran, J. C. M., Nelson, A. D., Skirycz, A. “Identification and characterization of the heat-induced plastidial stress granules reveal new insight into Arabidopsis stress response.” Frontiers in plant science, 2020.

Christodoulou, D., Kuehne, A., Estermann, A., Fuhrer, T., Lang, P., Sauer, U. “Reserve flux capacity in the pentose phosphate pathway by NADPH binding is conserved across kingdoms.” Iscience, 2019: 1133–1144.

Diether, M., Nikolaev, Y., Allain, F. H., Sauer, U. “Systematic mapping of proteinLmetabolite interactions in central metabolism of Escherichia coli.” Molecular systems biology, 2019.

Diether, M., Sauer, U. “Towards detecting regulatory protein–metabolite interactions.” Current opinion in microbiology, 2017: 16–23.

Gallego, O., Betts, M. J., Gvozdenovic◻Jeremic, J., Maeda, K., Matetzki, C., Aguilar◻Gurrieri, C.,…, Gavin, A. C. “A systematic screen for protein–lipid interactions in Saccharomyces cerevisiae..” Molecular systems biology, 2010.

Gene Ontology Consortium. “The gene ontology project in 2008.” Nucleic acids research, 2008.

Giavalisco, P., Li, Y., Matthes, A., Eckhardt, A., Hubberten, H. M., Hesse, H.,…, Willmitzer L. “Elemental formula annotation of polar and lipophilic metabolites using 13C, 15N and 34S isotope labelling, in combination with high□resolution mass spectrometry.” The Plant Journal, 2011: 364–376.

Hastie, T., Tibshirani, R., Friedman, J. The elements of statistical learning: data mining, inference, and prediction. Springer Science & Business Media, 2009.

Huerta-Cepas, J., Szklarczyk,D., Forslund,K., Cook,H., Heller,D.,Walter,M. C. et al. “eggNOG 4.5: a hierachical orthology framework with improved functional annotations for eukaryotic, prokaryotic and viral sequences.” Nucleic acids research, 2016.

Jackson, E. K. “Discovery and roles of 2’, 3’-cAMP in biological systems.” Non-canonical Cyclic Nucleotides, 2015: 229–252.

Kosmacz, M., Gorka, M., Schmidt, S., Luzarowski, M., Moreno, J. C., Szlachetko, J.,…, Skirycz, A. “Protein and metabolite composition of Arabidopsis stress granules..” New Phytologist, 2019: 1420–1433.

Kosmacz, M., Luzarowski, M., Kerber, O., Leniak, E., Gutiérrez-Beltrán, E., Moreno, J. C.,…, Skirycz, A. “Interaction of 2’, 3’-cAMP with Rbp47b plays a role in stress granule formation.” Plant physiology, 2018: 411–421.

Kosmacz, M., Sokołowska, E. M., Bouzaa, S., Skirycz, A. “Towards a functional understanding of the plant metabolome.” Current opinion in plant biology, 2020: 47–51.

Lempp, M., Farke, N., Kuntz, M., Freibert, S. A., Lill, R., Link, H. “Systematic identification of metabolites controlling gene expression in E. coli.” Nature communications, 2019: 1–9.

Li, X., Gianoulis, T. A., Yip, K. Y., Gerstein, M., Snyder, M. “Extensive in vivo metabolite-protein interactions revealed by large-scale systematic analyses.” Cell, 2010: 639–650.

Li, X., Snyder, M. “Metabolites as global regulators: A new view of protein regulation.” Bioessays, 2011.

Link, H., Kochanowski, K., Sauer, U. “Systematic identification of allosteric protein-metabolite interactions that control enzyme activity in vivo.” Nature biotechnology, 2013: 357–361.

Luzarowski, M., Skirycz, A. “Emerging strategies for the identification of protein–metabolite interactions..” Journal of experimental botany, 2019: 4605–4618.

Luzarowski, M., Vicente, R., Kiselev, A., Wagner, M., Schlossarek, D., Erban, A.,…, Skirycz, A. “Global mapping of protein–metabolite interactions in Saccharomyces cerevisiae reveals that Ser-Leu dipeptide regulates phosphoglycerate kinase activity..” 2021: 1–15.

Mallam, A. L., Sae-Lee, W., Schaub, J. M., Tu, F., Battenhouse, A., Jang, Y. J.,…, Drew, K. “Systematic discovery of endogenous human ribonucleoprotein complexes.” Cell reports, 2019: 1351–1368.

McWhite, C. D., Papoulas, O., Drew, K., Cox, R. M., June, V., Dong, O. X.,…, Marcotte, E. L. “A pan-plant protein complex map reveals deep conservation and novel assemblies.” Cell, 2020: 460–474.

Moreno, C. J., Rojas, B. E., Vicente, R., Gorka, M., Matz, T.,…, Skirycz, A. “Tyr-Asp inhibition of glyceraldehyde 3-phosphate dehydrogenase affects plant redox metabolism (accepted).” EMBO J, 2021.

Pedregosa, F., Varoquaux, G., Gramfort, A., Michel, V., Thirion, B., Grisel, O.,…, Vanderplas, J. “Scikit-learn: Machine Learning in Python.” the Journal of machine Learning research, 2011: 2825–2830.

Perez-Riverol, Y., Csordas, A., Bai, J., Bernal-Llinares, M., Hewapathirana, S.,…, Eisenacher, M. “The PRIDE database and related tools and resources in 2019: improving support for quantification data.” Nucleic Acids Res, 2019.

Piazza, I., Kochanowski, K., Cappelletti, V., Fuhrer, T., Noor, E., Sauer, U., Picotti, P. “A map of protein-metabolite interactions reveals principles of chemical communication..” Cell, 2018: 358–372.

Protter, D. S., Parker, .. “Principles and properties of stress granules.” Trends in cell biology, 2016: 668–679.

Razaghi-Moghadam, Z., Sokolowska, E. M., Sowa, M. A., Skirycz, A., & Nikoloski, Z. “Combination of network and molecule structure accurately predicts competitive inhibitory interactions.” Computational and Structural Biotechnology Journal, 2021: 2170–2178.

Reznik, E., Christodoulou, D., Goldford, J. E., Briars, E., Sauer, U., Segrè, D., Noor, E. “Genome-scale architecture of small molecule regulatory networks and the fundamental trade-off between regulation and enzymatic activity..” Cell reports, 2017: 2666–2677.

Sokolowska, E. M., Schlossarek, D., Luzarowski, M., Skirycz, A. “PROMIS: Global Analysis of PROtein-Metabolite Interactions.” Current protocols in plant biology, 2019.

Szklarczyk, D., Morris, J. H., Cook, H., Kuhn, M., Wyder, S., Simonovic, M. et al. “The STRING database in 2017: quality-controlled protein–protein association networks, made broadly accessible.” Nucleic acids research, 2016.

Szklarczyk, D., Santos, A., von Mering, C., Jensen, L. J., Bork, P., Kuhn, M. “STITCH 5: augmenting protein-chemical interaction networks with tissue and affinity data.” Nucleic Acids Research, 2016.

Thirumalaikumar, V. P., Wagner, M., Balazadeh, S., Skirycz, A. “Autophagy is responsible for the accumulation of proteogenic dipeptides in response to heat stress in Arabidopsis thaliana..” The FEBS journal, 2021: 281–292.

Veyel, D., Kierszniowska, S., Kosmacz, M., Sokolowska, E. M., Michaelis, A., Luzarowski, M.,…, Skirycz, A. “System-wide detection of protein-small molecule complexes suggests extensive metabolite regulation in plants..” Scientific reports, 2017: 1–8.

Veyel, D., Sokolowska, E. M., Moreno, J. C., Kierszniowska, S., Cichon, J., Wojciechowska, I.,…, Skirycz, A. “PROMIS, global analysis of PROtein–metabolite interactions using size separation in Arabidopsis thaliana..” Journal of Biological Chemistry, 2018: 12440–12453.

Wan, C., Liu, J., Fong, V., Lugowski, A., Stoilova, S., Bethune-Waddell, D.,…, Emili, A. “ComplexQuant: high-throughput computational pipeline for the global quantitative analysis of endogenous soluble protein complexes using high resolution protein HPLC and precision label-free LC/MS/MS..” Journal of proteomics, 2013: 102–111.

Wang, C. “Kernel learning for visual perception.” Doctoral dissertation, 2019.

Wang, Y., Suzek, T., Zhang, J., Wang, J., He, S., Cheng, T. et al. “PubChem bioassay: 2014 update.” Nucleic acids research, 2013.

Webb, E. C. “Enzyme nomenclature 1992. Recommendations of the Nomenclature Committee of the International Union of Biochemistry and Molecular Biology on the Nomenclature and Classification of Enzymes (No. Ed. 6).” Academic Press, 1992.

